# DNA-controlled Spatio-temporal Patterning of a Cytoskeletal Active Gel

**DOI:** 10.1101/2021.08.02.454703

**Authors:** Yuliia Vyborna, Jean-Christophe Galas, André Estevez-Torres

## Abstract

Living cells move and change their shape because signaling chemical reactions modify the state of their cytoskeleton; an active gel that converts chemical energy into mechanical forces. To create life-like materials, it is thus key to engineer chemical pathways that drive active gels. Here, we describe the preparation of DNA-responsive surfaces that control the activity of a cytoskeletal active gel comprised of microtubules: a DNA signal triggers the release of molecular motors from the surface into the gel bulk, generating forces that structure the gel. Depending on the DNA sequence and concentration, the gel forms a periodic band pattern or contracts globally. Finally, we show that the structuration of the active gel can be spatially controlled in the presence of a gradient of DNA concentration. We anticipate that such DNA-controlled active matter will contribute to the development of life-like materials with self-shaping properties.

---

Living cells and tissues change their shape autonomously because their cytoskeleton is an active gel that converts ATP chemical energy into mechanical work.^1^ To control their deformations during cell locomotion and development, cells and tissues tune the active state of their cytoskeleton over space and time. This is performed by chemical reactions that link the concentration of a trigger to the force exerted by the active gel.^2^ Despite its importance in the development of life-like synthetic materials,^3,4^ engineering such a chemo-mechanical trigger remains challenging. An important body of work has developed methods to chemically actuate hydrogels.^5,6^ However, current approaches either pattern passive hydrogels,^7–10^ or trigger deformations in active gels with chemical reactions that are difficult to design.^11,12^

A promising route relies on DNA hybridization reactions triggering forces that change the shape of hydrogels. Indeed, DNA reactions can be exquisitely controlled,^13–17^ and coupled to a great variety of materials.^18–22^ The mechanical amplification of a DNA signal can be either performed by DNA reactions, such as in DNA-responsive hydrogels,^23,24^ or by motor proteins, such as in DNA-based cytoskeletal active gels.^25–29^ While the former have the advantage of greater programmability, they lack the self-organization properties of cytoskeletal active gels.^30–33^ Despite progress in using DNA to tune the rheology of active gels^29,34^ or to trigger its activity at the micro-scale,^25,26,35^ we lack methods to control the spatio-temporal self-organization of cytoskeletal active gels at the macro-scale, as we demonstrate in the following.

The active gel used here is composed of clustered kinesin motors and microtubule filaments that are bundled together in the presence of a depletion agent. Because the filaments are continuously been propulsed by the motors such gels are known to self-organize into a variety of patterns.^31,36–40^ To control the activity of such gels with a DNA signal, kinesin clusters were attached to the DNA-modified surface of a glass channel filled with a solution of microtubule bundles (Fig. 1A).^41,42^ Motor clusters were released to the medium solely after the addition of DNA input **I**, thus initiating the continuous movement of microtubules and generating band structures followed by a global contraction of the gel (Fig. 1C). We demonstrate that the amplitude of these patterns is controlled by the sequence and the concentration of DNA. Furthermore, when the input DNA strand was heterogeneously distributed, it acted as a morphogen and provided a spatial control over the patterns observed in the active gel. Thus, this system paves the way toward the on-demand shaping of soft matter guided by DNA.

**Figure 1.**
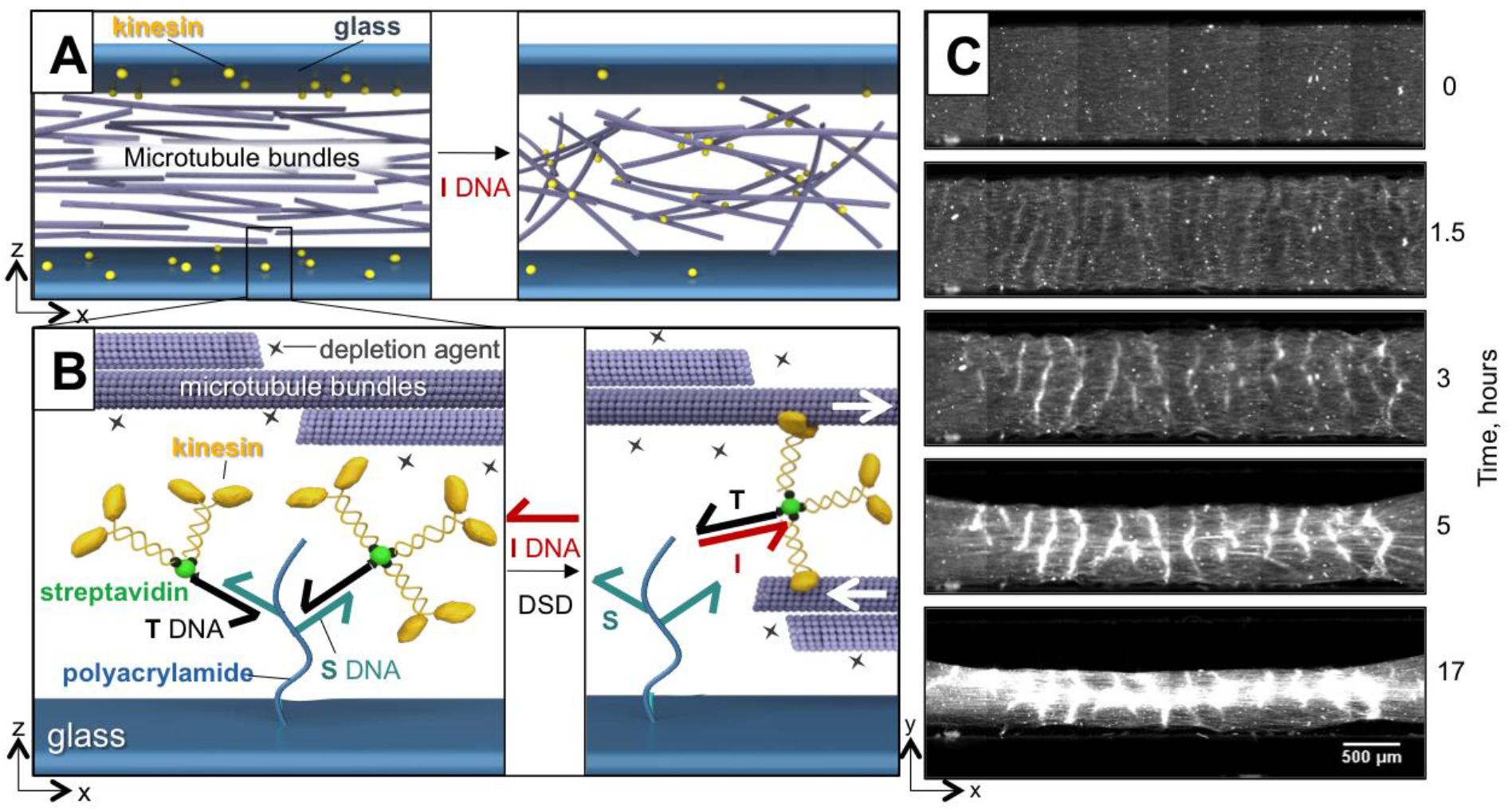
DNA-responsive, motor-decorated surfaces control the patterning of a cytoskeletal active gel. A: Schematic representation of a channel filled with microtubule bundles. Glass surfaces (in blue) are modified with kinesin-DNA constructs (yellow spheres). Upon addition of strand **I**, the motors are released into the bulk inducing forces upon the microtubules. B: Schematic representation of a glass surface functionalized by DNA/polyacrylamide brushes with kinesin–streptavidin-DNA constructs attached through DNA hybridized strands that undergo toehold-mediated strand displacement reaction. An invading strand **I** binds and displaces **T** DNA, releasing kinesin-streptavidin-DNA constructs that bind to microtubules and move them relative to each other (white arrows). Microtubules form bundles due to the attraction interactions induced by depletion agent Pluronic (black stars). C: Time-lapse fluorescent microscopy images of the microtubule solution confined in glass channels in the presence of 1 μM of **I** DNA. For detailed conditions see SI.

To prepare densely-covered DNA glass surfaces, acrylamide and acrydite-DNA **S** were copolymerized onto silanized glass slides (SI).^43^ These DNA-modified slides were further used to fabricate rectangular channels with typical dimensions of 22 × 1.5 × 0.1 mm^3^. Then, a solution of biotin-labeled target DNA **T** was added to the channel to form duplex **S**:**T** on the surface due to their partial complementarity (Fig. 1B). Subsequently, streptavidin (**str**) was injected into the channel and conjugated with biotinylated **T**, generating complex **S**:**T**:**str**. Considering that **T** bears a single biotin and that streptavidin was added in 10-fold excess, the formation of 1:1 **T**:**str** complexes with three free biotin-binding sites in the streptavidin was expected. Finally, biotinylated kinesin (**kin**) was flown inside the channel to form **S**:**T**:**str**:**kin_n_** with a **kin**/**str** ratio, n, that varied between 1 and 3. This resulted in kinesin clusters attached to the surface and distributed homogeneously throughout the channel. Fluorescently labeled streptavidin was used to estimate the surface density of motor clusters that could be changed from 3 × 10^8^ to 2 × 10^10^ molecules/cm^2^ (SI Fig. S1). The non-specific attachment of **T**, **str** and **kin** to the surface was minimized in the presence of a 2% solution of Pluronic F-127 (SI Fig. S2).

The kinesin-decorated channel was subsequently filled with a solution of microtubules (Fig. 1A) self-assembled into bundles in the presence of the depletion agent Pluronic.^31,44^ The detachment of kinesin clusters from the surface was achieved through a DNA strand displacement (DSD) reaction in the presence of the invader strand **I** that is fully complementary to **T** (Fig. 1B).^45^ We thus have

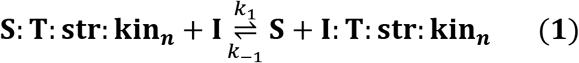

where the formation of the duplex **I**:**T** is thermodynamically favored compared to **S**:**T** because the former is 5 base-pairs longer than the latter. Fig. 1C and Movie S1 show the behavior of the gel in the presence of 1 μM of **I**. In the first 3 hours, we observed the formation of bright bands parallel to the *y* axis with a typical period of 300 ± 50 μm. These bands are reminiscent of the corrugations observed in a similar system where the kinesin clusters were initially present in solution.^37^ Afterwards, the gel contracted along *y* and *x*, making the bands brighter. Noteworthy, no bands or contraction was observed in the absence of **I** or in the presence of a DNA strand with a random sequence (**N**) (Fig. S3 and Movie S1). The physical separation between the motors and the microtubules was key to obtain a robust inactive initial state. In a control experiment with kinesin clusters present in solution from the start, bands and contraction were also observed (Fig. S11). These observations are compatible with reaction (1), indicating that **I** releases kinesin clusters into the medium, inducing the mutual sliding of the bundles and shaping the active gel at the centimeter scale.

We measured the kinetics and thermodynamics of the forward reaction (1) by following the fluorescence from labeled-streptavidin on the glass surfaces. Fig. 2A shows that the characteristic time for releasing **I**:**T**:**str**:**kinn** from the surface is equal to 5.5 min when [**I**] = 1 μM and thus the corresponding rate constant is *k*_1_ = 3 × 10^3^ M^−1^s^−1^, which is compatible with previous measurements of dsDNA association kinetics on surfaces^46^ and 10^3^ times smaller than the typical value in solution.^47^ In the absence of **I**, the streptavidin-associated fluorescence remained unchanged over time. To quantify active gel dynamics we defined a gel contraction index as the standard deviation of the microtubule intensity (see SI Section 6). The dynamics of the gel was significantly slower compared to the kinetics of the DNA reaction (Fig. 2B), with a characteristic time of 300 min. Fig. 2C (red) shows the titration of kinesins attached to the surface by increasing concentrations of **I**. We extracted an equilibrium constant K1 = 2.3 × 10^−2^ for reaction (1), similar to the one measured in the absence of microtubules (K*_1_ = 3.6×10^−2^, SI Fig. S4) confirming that reaction (1) is driven by DNA hybridization and not by a potential kinesin-microtubule interaction. We estimated the corresponding equilibrium constant in solution at 22 °C to be 4 × 10^6^ (NUPACK) indicating that reaction (1) is significantly hindered on the surface compared to solution, as expected.^48^ Importantly, increasing [**I**] resulted in an increasingly contracted microtubule network (Fig 2C black) and we observed network structures from slightly contracted at 5 nM, to bright bands at 50 nM and strong global contractions at ≥ 250 nM (SI Fig. S7).

**Figure 2:**
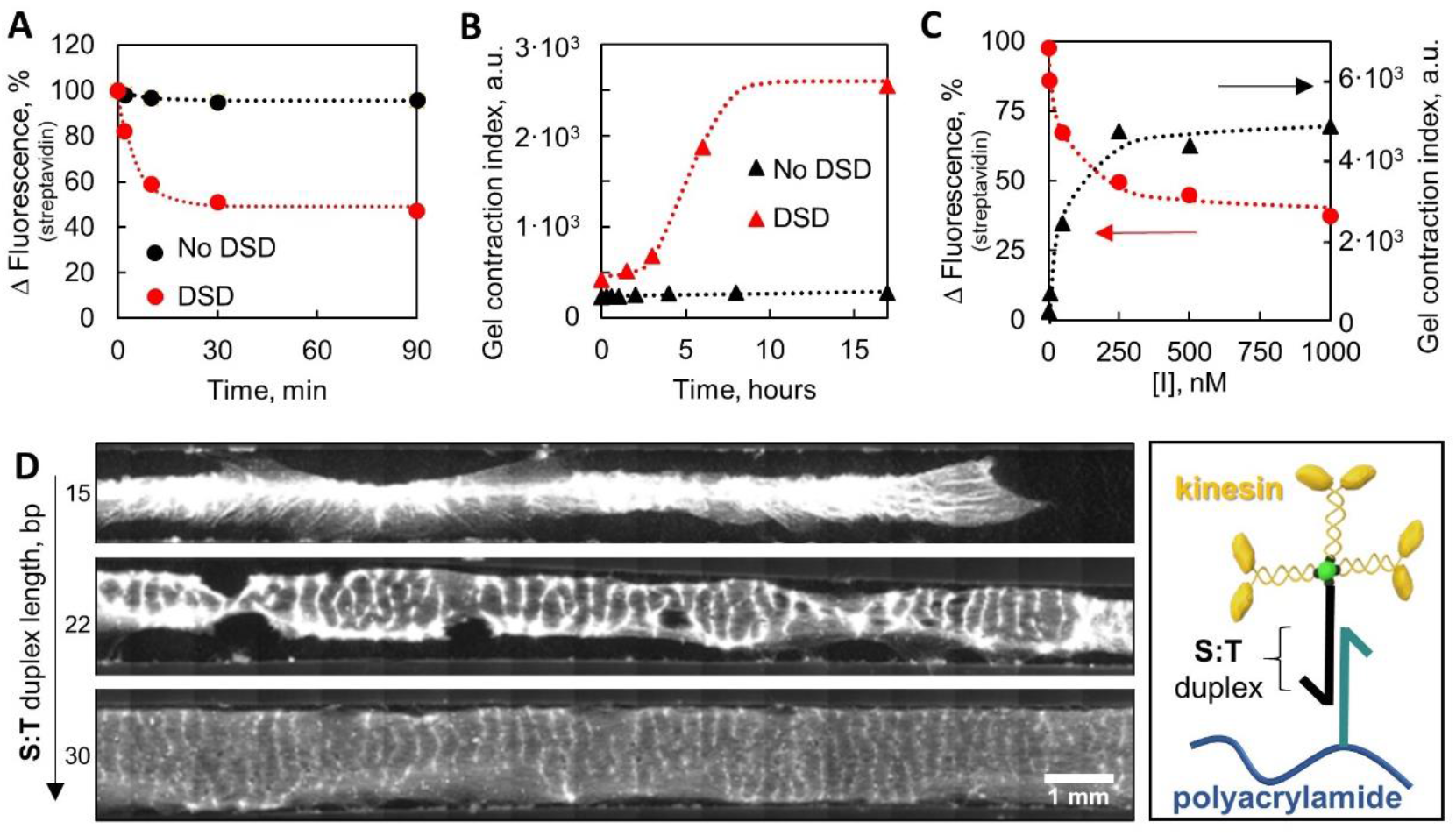
Influence of the kinetics and thermodynamics of the DSD reaction on the active gel patterning. Streptavidin fluorescence on the surface of the glass (A) and gel contraction index (B) as a function of time in the absence and in the presence of 1 μM of **I**. The dotted lines are fits to eqs. S2 and S8 respectively. Streptavidin fluorescence on the surface of the glass after 90 minutes (red) and gel contraction index after 17 hours (black) as a function of the concentration of **I** (C). The dotted lines are fits to eqs. S6 and S7. D: Fluorescence microscopy images of microtubules confined in glass channels modified with biotinylated DNA strands of different complementarity lengths in the presence of 1 μM **I** strand after 18 hours. For detailed conditions see SI.

Changing the length of the duplex in the complex **S**:**T**, which allows attaching kinesin clusters to the surface, provides another way to control the activity of the gel via the DSD reaction. Indeed, shorter **S**:**T** duplexes are expected to increase K1 and thus the final concentration of kinesin clusters in solution. In agreement with this interpretation, the amplitude of the gel deformations increased as the length of the duplex decreased at constant [**I**] = 1 μM (Fig. 2D and Movie S2). We observed faint regular bands for the 30 bp-long duplex, bright bands with local gel breaks for 22 bp, and bright bands with global contraction for the 15 bp-long duplex. Confocal images showed that faint bands correspond to a local increase in the microtubule density while bright bands were associated with gel corrugations in the *yz* plane (Movie S4). A third parameter that allows to control gel activity through DSD reaction is the concentration of the immobilized kinesin on the surface. Fig. S8 examines the effect of **str** surface density on the dynamics of the active gel for [**I**] = 1 μM. For the densities of 0.5 ×, 1 × and 2 × 10^10^ molecules/cm^2^ we respectively observed: slight bulk contraction with no bands, weak bands and bright bands with strong global contraction. Taken together, the data from Fig. 2, S3 and S7–S8 imply that the DSD reaction is fast compared to the dynamics of the active gel and that the state of the gel is determined by the concentration of motors released in solution, which is tuned by the thermodynamics of the DSD reaction and by the surface density of **str**.

In biology, morphogens play a crucial role in the patterning of developing tissues by diffusing in space and forming concentration gradients that provide positional information to downstream molecular events. In particular, morphogen gradients influence active matter processes in vivo.^49^ In the following, we assessed the invader DNA strand as a morphogen that provides positional information to an active gel. To do so, 1μM of DNA **I** (the morphogen) was put solely on the left side of a channel filled with microtubules (Fig. 3 and SI Methods). On this side of the channel the onset of gel activity was detected after two hours followed by the formation of bright spots and global contraction in the *y* direction. As expected, **I** diffused toward the right side forming a concentration gradient of **I**, and thus of released kinesin clusters, that contracted the gel with decreasing amplitude and increased delay as the distance from the injection point increased (Fig. 3, Fig. S9 and Movie S3). In the central part of the channel, **I** arrived by diffusion at a concentration significantly lower than 1 μM, and activity was observed after 10 hours with a front of activity progressing from left to right, with bright bands appearing sequentially. No activity was observed over the course of 43 hours in the right side of the channel where the morphogen concentration was negligible.

**Figure 3:**
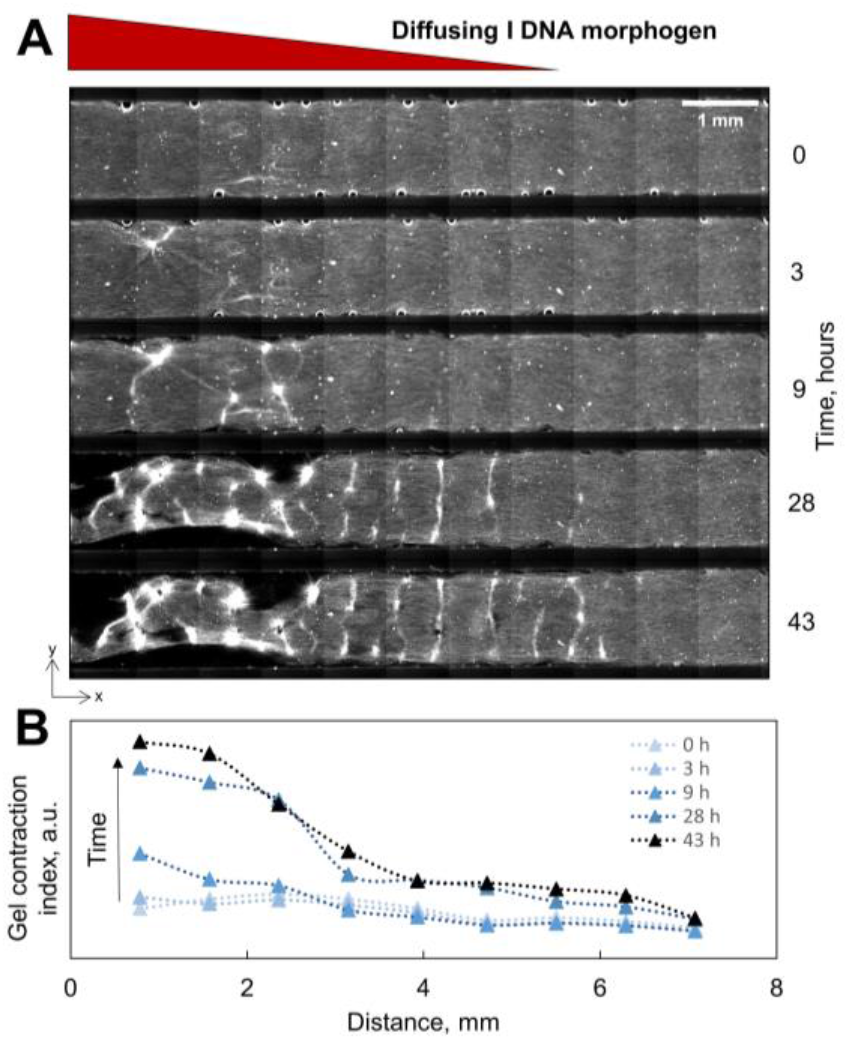
Active gel patterning through an underlying morphogen gradient of DNA. A: Time-lapse fluorescent microscopy images of the microtubule solution confined in glass channels after the addition of 1 μM **I** DNA on the left side. For detailed conditions see SI. B: Gel contraction index computed along the channel length for each time point (Fig. S6). The dotted lines are guides to the eye.

The multiscale structuring of shapeless soft matter, including active materials, is conceptually and experimentally challenging.^5^ Here it was shown that the large-scale mechanical self-organization of microtubules could be activated in a programmable manner by a DNA signal. DNA-kinesin conjugates hybridized to a DNA-modified surface could be released in a controllable way by specific DNA inputs that trigger a DNA strand displacement reaction. Moreover, the DNA input could be distributed heterogeneously across space and act as a morphogen, transforming chemical information into a mechanical and structural output with spatial resolution. Complementary to the systems that utilize mechano-chemical transduction,^50^ this work provides an approach allowing for preparation of a synthetic material that actuates along a chemo-mechanical pathway. We anticipate that this methodology will open a new route to accessing novel self-fabricated, force-exerting synthetic soft matter with the potential of integration in soft robotics and biological environments.

## SUPPORTING INFORMATION

### 1. Methods

#### 1.1. Oligonucleotides

All oligonucleotides used in this study were purchased HPLC-purified from biomers.net GmbH. The immobilized **S** strand is 35-nt long. DNA targets **T-30**, **T** and **T-15** have, respectively, sequences complementary to 30, 22, or 15 bases of **S**, with a 5-nt overhang at the 5’ end. The invader **I** is fully complementary to **T-30**, both 35-nt long. Non-specific binding of the target DNAs was checked by using biotinylated non-complementary DNA **NT**. Non-specific DSD reaction was checked with fluorescently-labeled non-complementary DNA **N**. The sequences of the DNA are listed in Table S1.

**Table S1.**
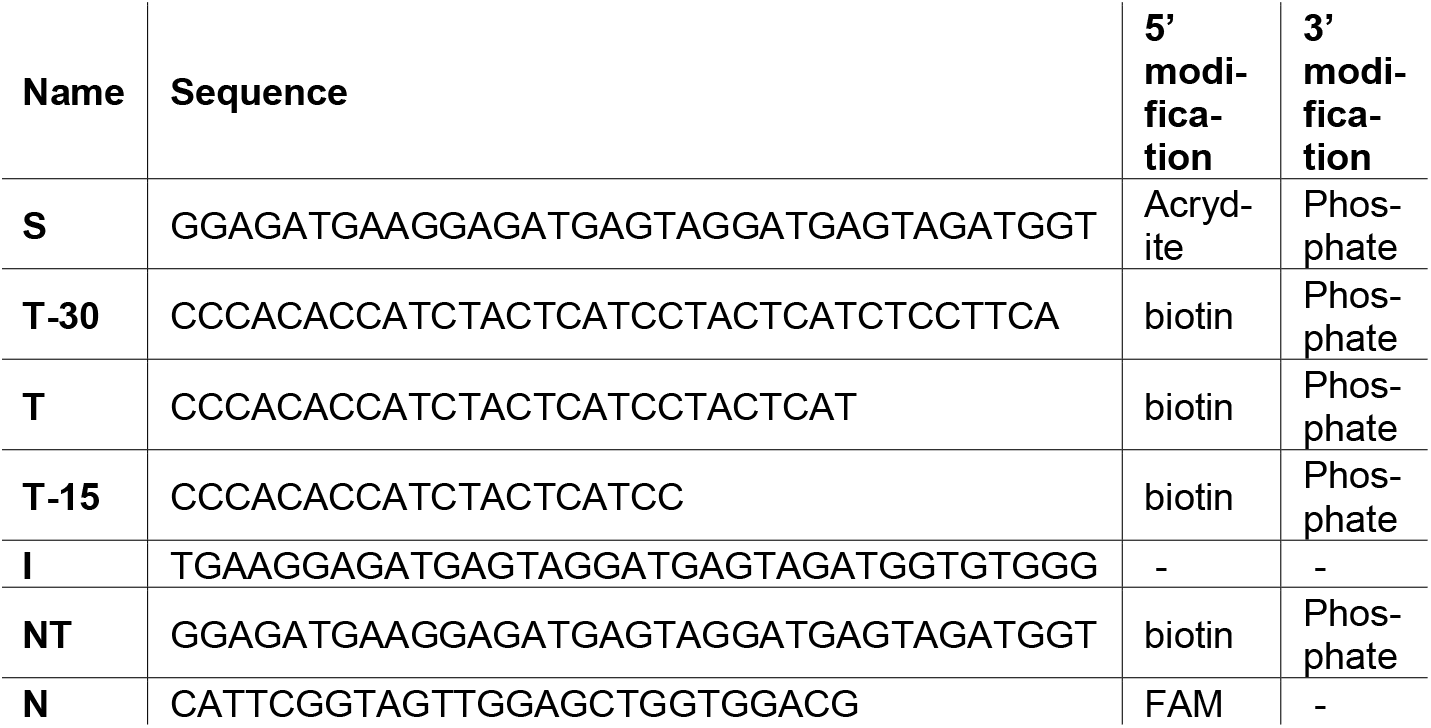
DNA sequences used in the study.

#### 1.2. Kinesin and microtubules preparation

Biotin-labeled kinesin K401 was purified as described in ref. 1. Purified tubulin and rhodamine-labeled tubulin protein for microtubules preparation were bought from Cytoskeleton Inc. Taxol-stabilized microtubules were prepared as described in ref. 2 and were incubated at room temperature for at least 24 hours before using them in the experiments.

#### 1.3. Glass modification

To prepare substrates for DNA immobilization, glass coverslips were first cleaned by sonication over 10 minutes in 0.5 % Hellmanex (Sigma), the same procedure was repeted in ethanol and finally, in 0.1 M KOH.^3^ The slides were subsequently rinsed in water. Next, the glass slides were treated with silane-coupling solution (1% acetic acid and 0.5% 3-(Trimethoxysilyl)propylmethacrylate in ethanol) for 15 minutes and rinsed in water. 200 μl of an aqueous solution containing 2% acrylamide, 0.02% 5′-acrylamide oligonucleotide **S**, 0.125% w/v APS and 0.125% v/v TEMED was manually pipetted onto silanized slides. The solution was covered with a second silanized coverslip, making a sandwich, and placed in a water-saturated, argon atmosphere inside a desiccator; at room temperature for 3 hours to allow polymerization. Non-immobilized DNA and polyacrylamide were removed from the surface by washing in water, then air-dried and stored at room temperature. The calculated maximum surface density of **S** in this DNA-modified glass slide was 7·10^13^ mol/cm^2^.

#### 1.4. Channel assembly

Channels were assembled using a DNA-modified glass slide (26 × 75 × 1 mm) and a DNA-modified coverslip (22 × 60 × 0.17 mm) separated by Parafilm cut with a Graphtec Cutting Plotter CE6000-40, gently melted over a heating plate. The 22×1.5×0.1 mm^3^ fabricated channel had a volume of ~ 3.5 μL and a coated surface of ~ 0.7 cm^2^. The thickness of the channel may vary from 0.11 to 0.9 mm. Thus, the resulting amount of the tubuline is equal to 2.15±0.15×10^13^ molecules/cm^3^. This variation did not exhibit any significant difference in the experiments.

#### 1.5. Kinesin conjugation to the glass surface of the channel

Before each experiment, channels were treated with a 2% solution of Pluronic127 in 1x PBS at room temperature in a humid chamber for one hour to block unspecific protein adsorption and subsequently washed with 50 μl of 1x PBS (in both directions). To create a final concentration of kinesin clusters on the surface equal to 1×10^10^ molecules/cm^2^, 10 μl of a 20 nM solution of biotin-labeled complementary DNA (**T**, **T-15**, or **T-30**) in 1x PBS was flushed through each channel and allowed to hybridize to the surfaces for one hour at 65 °C in a humid chamber placed on the heating plate. After the subsequent washing with 50 μl of 1x PBS (in both directions), 10 μl of the 20 nM solution of Atto 488-tagged streptavidin **str** (Sigma-Aldrich) was added and incubated at RT for 10 min. Non-conjugated streptavidin was washed with 50 μl of 1x PBS (in both directions). Lastly, 10 μl of 50 nM of biotin-labeled kinesin (K401) **kin** in 1x PBS was added to each channel and incubated for 15 min at RT to conjugate to the surface-immobilized streptavidin/DNA constructs. The excess of the kinesin was washed with 50 μl of 1x PBS (in both directions), and the channels were used immediately for further experiments.

#### 1.6. Active gel experiments

After the kinesin conjugation, 4 μl of the microtubules solution with the required concentration of the components, as indicated in Table S2, was filled in the channel by capillarity, sealed with vacuum grease and placed on the microscope stage. In most of our experiments we used a Tokai-Hit temperature control microscope stage set at 22 °C in a room with air conditioning set at 21 ± 1°C. In some occasions, the temperature-controlled stage was not used. We did not observe differences between the former and the latter experiments. When the temperature stage controller was used, mineral oil was added between the glass channels and the stage.

#### 1.7. Kinetics and thermodynamics experiments in Fig. 2A and Fig. 2C (red)

To track the changes of the fluorescence of the streptavidin on the surface, the channels were filled with solutions (Table S2) and were left open to enable subsequent washing. After the incubation time indicated in Table S2, the channels were washed with 50 μl of 1x PBS and the fluorescence was recorded.

#### 1.8. Gradient experiment (Fig. 3)

To create a non-homogeneous concentration profile of **I** DNA inside the channel, and thus mimic the morphogen gradients that provide positional information to downstream molecular events in biology, we used a micro-pipette to first fill ~75% of the length of the functionalized channel with 2.25 μl of a solution of microtubules without **I.** Then we filled the remainder ~25% of the channel (0.75 μl) with a solution of microtubules containing 1 μM **I** DNA. Once sealed, the active gel was imaged with fluorescence micro-copy. To monitor at the same time the diffusion of **I**, we added a fluorescent DNA strand of similar length**, N** (Fig. S9).

#### 1.9. Imaging

Epifluorescence images of the channels were obtained at 22°C with a Zeiss Observer 7 automated microscope equipped with a Hamamatsu C9100-02 camera, a 10× objective, and a motorized stage controlled with MicroManager 1.42. Images were recorded with excitations at 470 or 550 nm with a CoolLED pE2. ***Image Analysis*** was done with ImageJ.

**Table S2.**
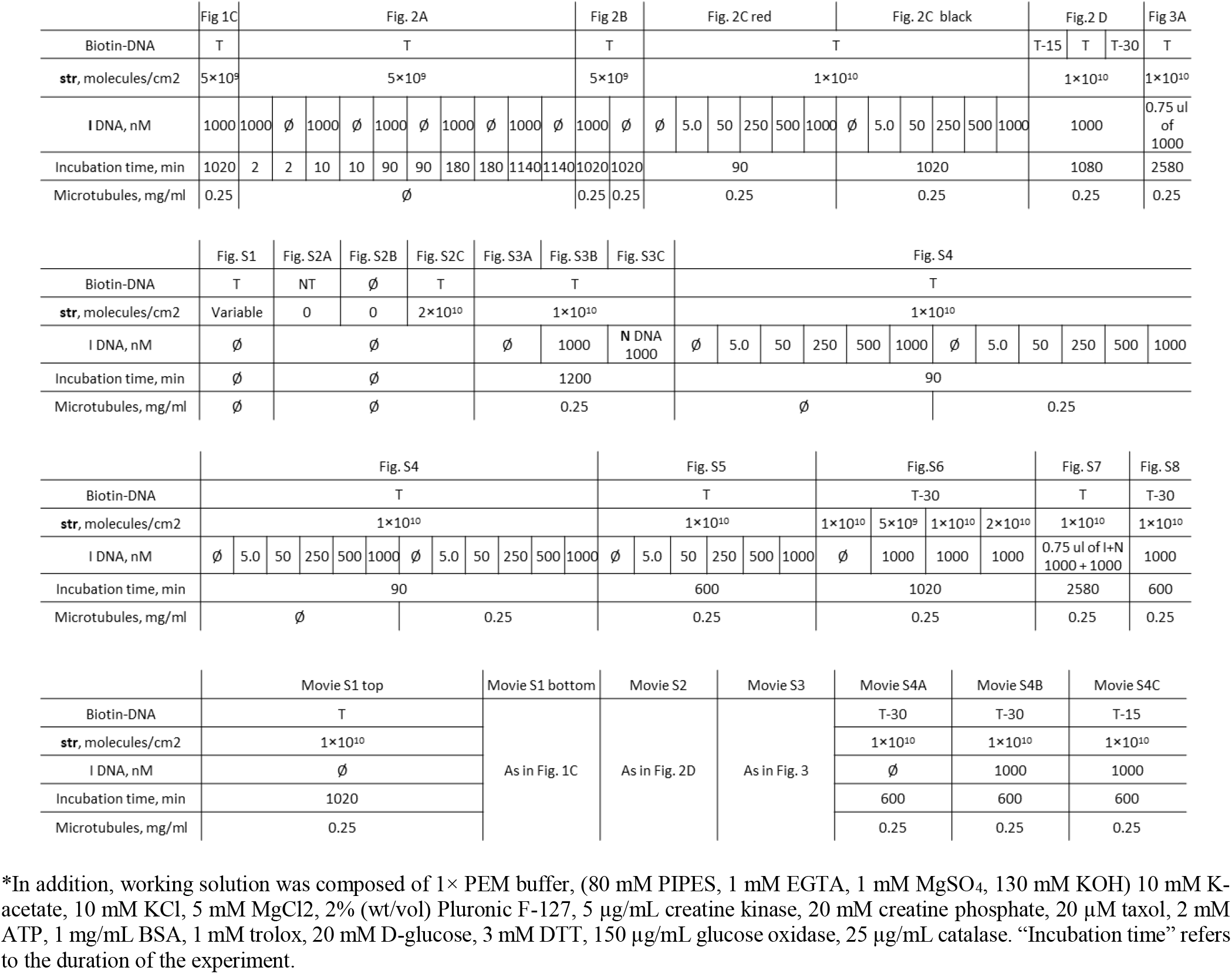
Summarized experimental conditions.*

### 2. Quantifying the density of the streptavidin on the surface

We estimated the average surface density of streptavidin for each experiment by using fluorescently-labeled **str**. Glass channels with hybridized biotin-tagged DNA were filled with the solutions of the fluo-rescently-labeled streptavidin with concentrations ranging from 0 to 50 μM and we recorded the fluorescence signal before the washing step from which we subtracted background fluorescence to obtain *F*_bw_. We obtained a linear relation that we fitted with *F*_bw_ = *a*[**str**]_bulk_. Then, the channels were washed with 50 μl PBS buffer and the fluorescence was recorded again, noted *F*_aw_. We then calculated the equivalent concentration of **str** on the surface as [**str**]_surf_ = (*F*_aw_)/*a*, and the corresponding surface density considering the volume (*V* = 3.5 μL) and the surface of the channel (*S* = 0.7 cm2) is ρ = (*F*_aw_)*V*N_A_/(*a*S)*. A control experiment with surfaces lacking the biotinylated **T** strand indicated that the non-specific binding of **str** to the surface was negligible.

**Figure S1.**
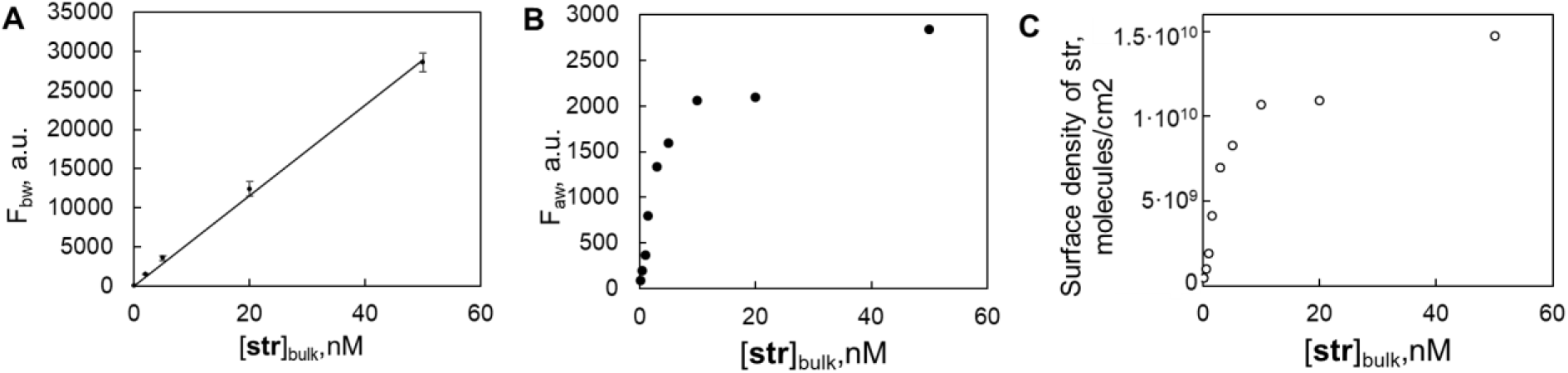
A: Fluorescence intensity before washing as a function of streptavidin concentration in bulk. The data points and error bars correspond, respectively, to the averages and standard deviations obtained from a triplicate. The line is a linear fit to the data. B: Fluorescence intensity after washing as a function of streptavidin concentration in bulk. C: Correlation between surface density of the streptavidin and concentration in the bulk.

### 3. Evaluation of the specific attachment of proteins and DNA to the surface

**Figure S2:**
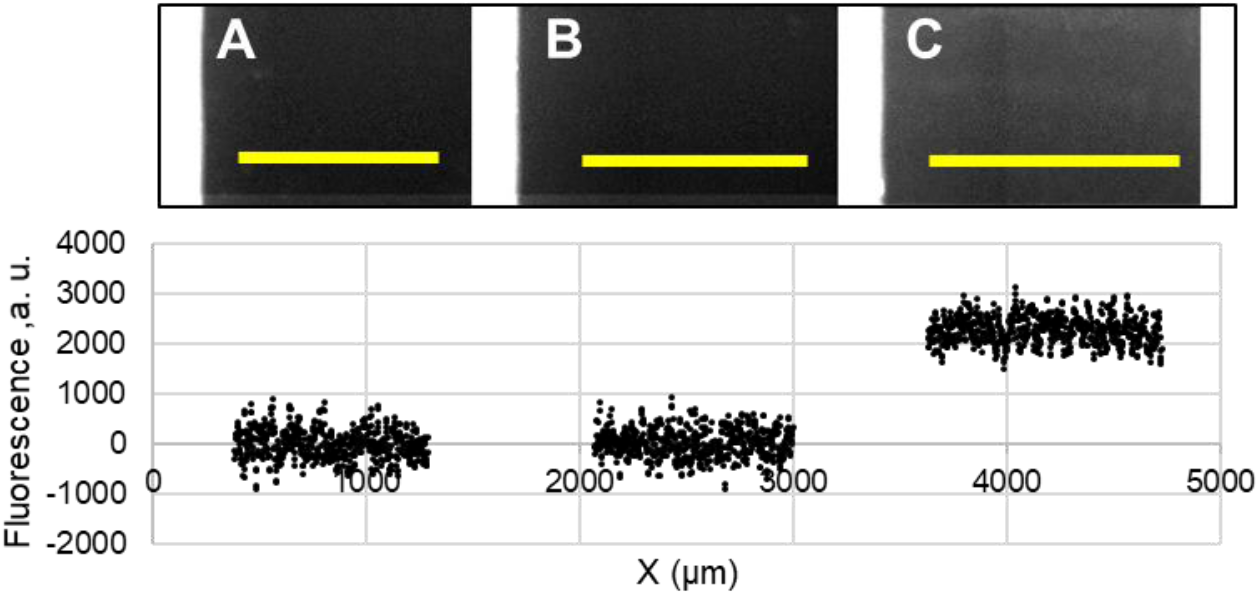
Top: Fluorescence images of the surfaces treated with 20 nM of non-complementary biotinylated **NT** DNA (A), no DNA (B), and 20 nM complementary biotinylated **T** DNA (C) and subsequently treated with 20 nM fluorescently-labeled streptavidin for 10 min and washed with buffer. Bottom: fluorescence intensity profiles along the yellow line depicted in the top figures. The results indicate a selective attachment between the **T** and the DNA-modified surface. Control surfaces did not show signs of non-specific adsorption of streptavidin or biotinylated DNA.

### 4. Control experiments of DNA strand displacement reaction showing the influence of invader complementarity and the presence of the microtubules

**Figure S3:**
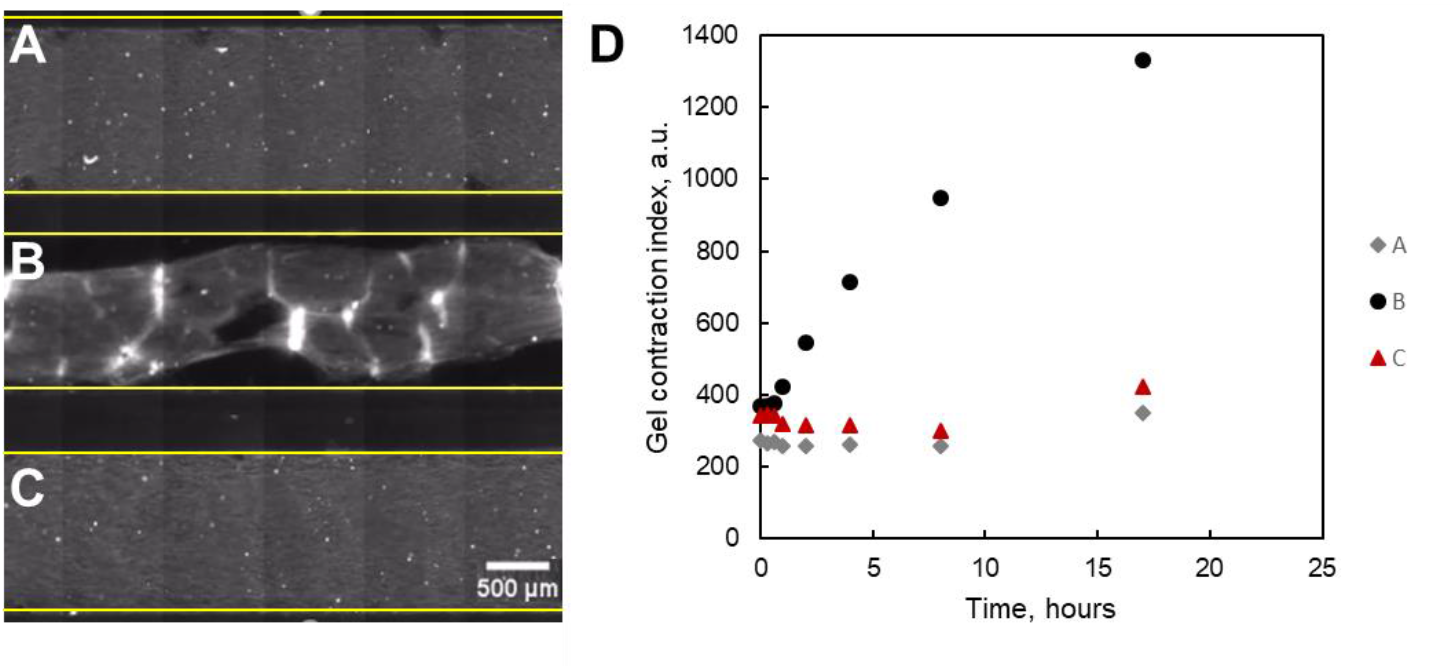
The kinesin-decorated surfaces are stable in the absence of complementary DNA **I**. Microtubule fluorescence images after 20 h for [**I**] = 0 (A), [**I**] = 1 μM (B) and [**N**] = 1 μM (C). No activity was observed in the presence of non-complementary DNA **N** (C). The borders of the channels are marked in yellow. D: Corresponding standard deviation over time of the microtubule fluorescence for the data described in panels A-C. For detailed conditions see Table S2.

**Figure S4:**
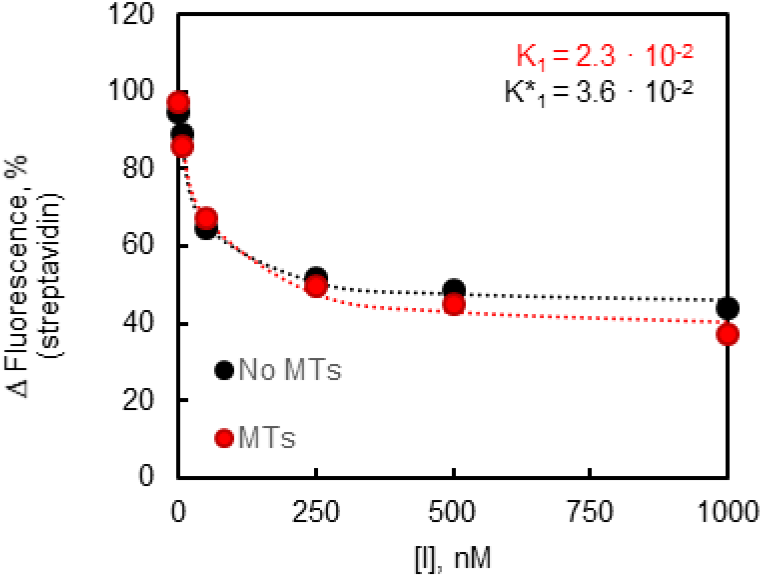
The release of **str** from the surface by **I** is independent of the presence of microtubules. **Str** fluorescence on the surface of the glass after 90 minutes for different concentrations of **I** in the presence (red) and in the absence (black) of microtubules. Equilibrium constants K_1_ and K*_1_ of reaction (S1) were found by fitting the data to the equation S6.

### 5. Models for the kinetics and thermodynamics of the DSD reaction and for the kinetics of gel activation

#### 5.1. Kinetics of the DSD reaction

We measured the detachment of the kinesin–streptavidin-DNA construct from the surface due to the DSD reaction in the presence of **I** by following the fluorescence intensity of **str** on the surface of the glass. We can write:

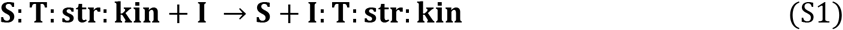

Because **I** is in excess relative to **S**:**T**:**str**:**kin**, we can consider reaction (S1) as pseudo-first-order with an apparent rate constant κ_1_ = *k*_1_[**I**]_0_, with [**I**]_0_ = 1 μM in our experiments. We note **Y** = **S**:**T**:**str**:**kin**, and thus

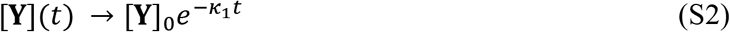

where the index 0 denotes the initial condition. In our experimental conditions, the change in fluorescence at the glass surface is expected to be *F(t)* = α[**Y**](*t*)+*F*_bk_, where α is a constant related to the fluorescence quantum yield of species **Y** that, in addition, integrates the intensity of the excitation light and the performance of the microscope optics and camera, and *F*_bk_ is the background fluorescence measured in the channel in the presence of buffer. As a result, to extract *κ*_1_, we normalize the fluorescence signal over time, *F(t)*, (data in Fig. 2A and S4) and fit it with:

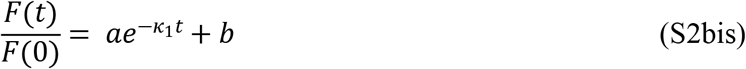

#### 5.2. Thermodynamics of the DSD reaction

We note **Z** = **I:T**:**str**:**kin** and thus reaction (1) in the Main Text now writes

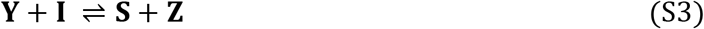

The equilibrium constant associated to the forward reaction is

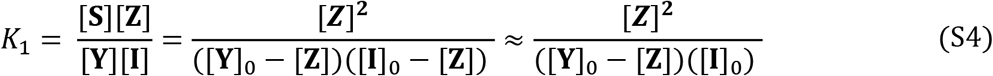

where the rightmost approximation is taken because species **I** is in excess. By solving the second-order polynomial, we find

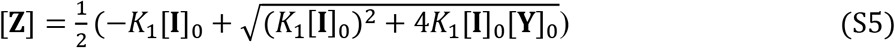

With [**Y**] = [**Y**]_0_-[**Z**] and the relation between [**Y**] and fluorescence *F* remaining valid, we have

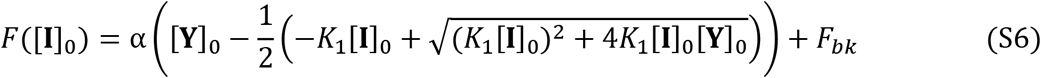

which we fitted to data of the **str** fluorescence on the surfaces as a function of [**I**] in (Fig. 2C red disks and Fig. S4) to extract *K*_1_.

Furthermore, if we suppose that that gel contraction index is proportional to [**Z**], the corresponding data as a function of [**I**] (Fig. 2C black triangles) was fitted to the equation:

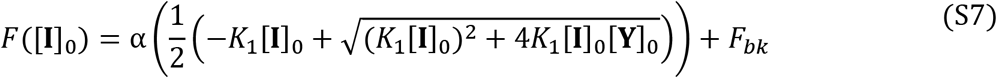

#### 5.3. Kinetics of gel activation

Fig. 2B in the Main Text shows the dynamics of the gel activation after the addition of DNA **I**. To extract a gel contraction rate, we heuristically fitted the data with the sigmoid equation

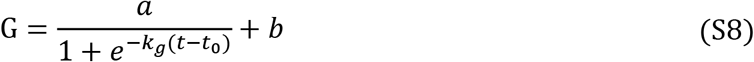

where G is the gel contraction index, calculated from the standard deviation of the fluorescence intensity of the microtubules, *k*_g_ is the rate constant associated to the contraction dynamics, *t*_0_ a delay associated to the saturation process of the contraction (due either to resource depletion or a force equilibration between active contraction forces and the passive rigidity of the gel) and *a* and *b* are constants associated to the initial and final level of the gel contraction index. For Fig. 2B, we obtained *k*_g_ = 60 min^−1^ and *t*_0_ = 300 min.

### 6. Characterization of the contraction of the active gels

When the gel contracts globally, the usual method to quantify the amplitude of contraction is to measure the area of the whole gel (i.e. the fluorescent area). Because we do not observe just a global contraction of the gel but rather an addition of local and global contractions and we want a measurement that is sensitive to both processes, we cannot use the area of the gel.

Alternatively, to measure the spreading state of the microtubules by taking into account both global contraction and local contractions, we measured the width of the histogram of pixel intensities in the channel. As depicted in Fig. S5 - associated to Fig. 2B in the main text – the contraction of the gel induces changes in these histograms. Initially, from t=0h to t=3h, in the absence of a global contraction, local contractions induce a widening of the intensity distribution, and the gel contraction index slightly increases from 415 to 678. From t=3h to t=6h, the local contractions are associated to a global contraction that induces the appearance on the histogram of a second sharp peak corresponding to the region of the channel devoid of microtubules. The gel contraction index then increases strongly from 678 to 1875. After 17h, the gel contraction index reaches 2549.

To measure the width of these histograms in a reproducible way, we first applied a Gaussian filter to the images, to blur them slightly, and then measured the standard deviation of the microtubule intensity, this value is what we call gel contraction index.

A local gel contraction index can also be obtained considering only a region of interest in the images (see Fig. S6).

**Figure S5:**
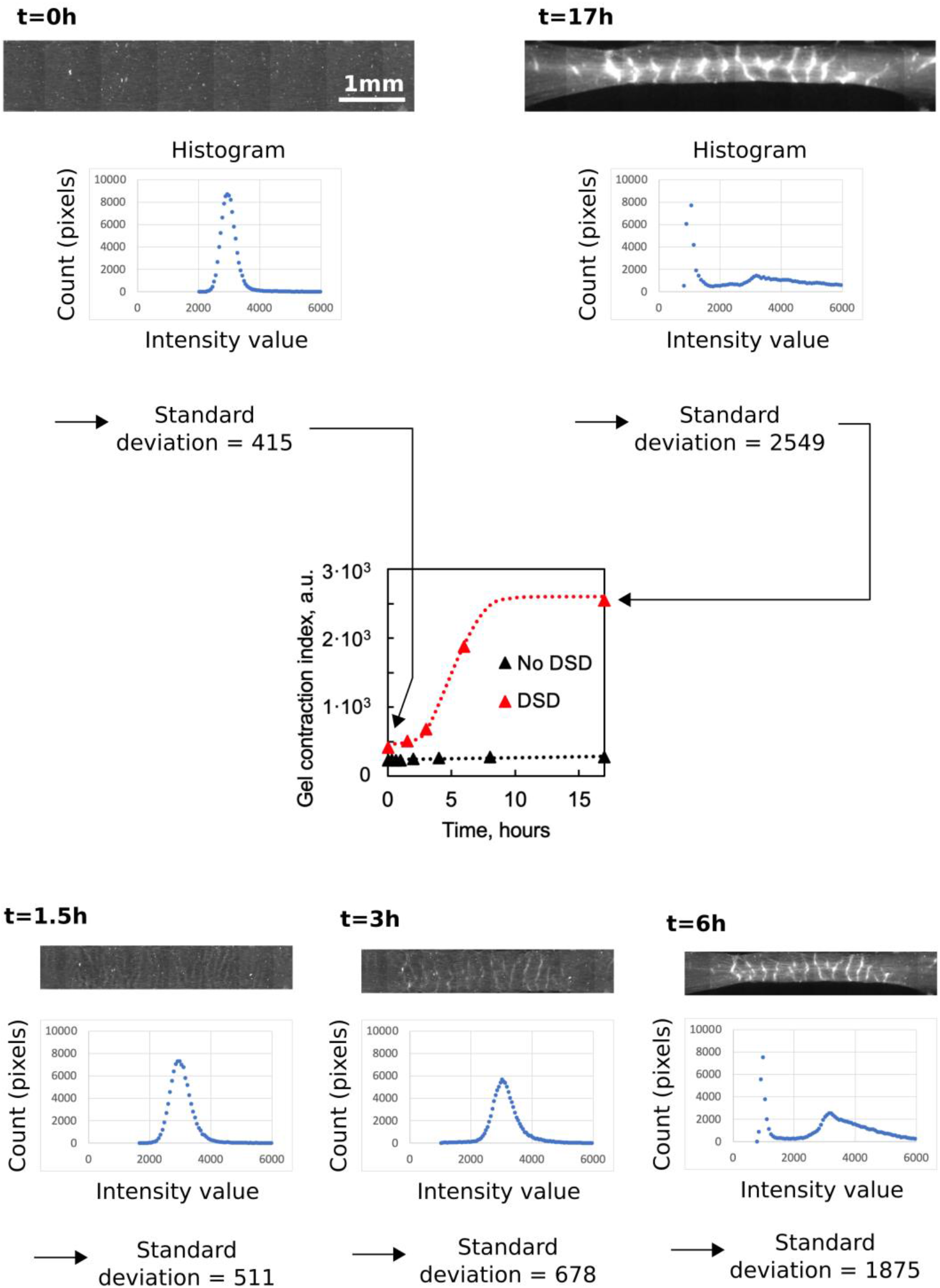
Contraction of an active gel over time. Fig. 2B from the main text is reproduced in the center and shows the gel contraction index as a function of time. For each timepoint of the contraction (t=0, 1.5, 3, 6 and 17 hours), after a gaussian filtering, the histogram of the imaged microtubules is plotted and used to extract the standard deviation of the microtubule intensity, which we define as a gel contraction index.

**Figure S6:**
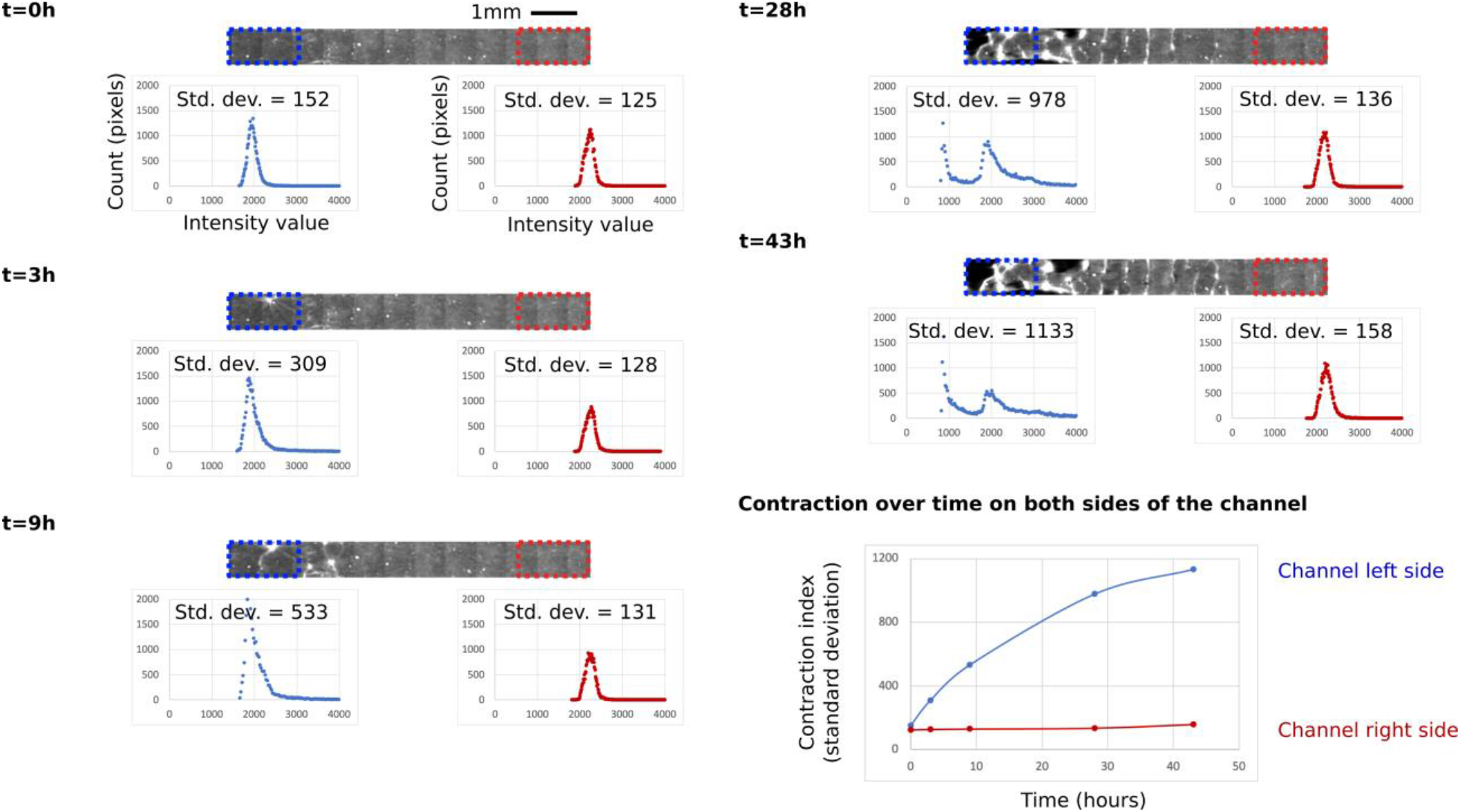
Inhomogeneous contraction of an active gel over time. This figure is associated to Fig. 3 in the main text which shows the inhomogeneous patterning of the active gel through an underlying gradient of DNA trigger. A local gel contraction index is determined on both sides of the channel considering ROI images, at different times (t=0, 3, 9, 28, 43 hours). At t=28 hours, the global contraction taking place on the left side of the channel induces the appearance on the corresponding histogram of a second sharp peak. The histogram of the imaged microtubules are used to extract the local standard deviation of the microtubule intensity, thus a local gel contraction index over space and time (bottom right panel).

### 7. Comparison of contraction rate with diffusion of DNA from the surface into the bulk

The release of motors from the surface into the bulk happens in two steps: the kinesin detaches from the surface via reaction (1) in MT and then it diffuses along the height of the channel within the gel, which ultimately causes its contraction. We measured the kinetics of the reaction step in Fig. 2A and demonstrate that its timescale is 5.5 min. We can estimate the diffusion time τ_D_ of kinesin clusters with typical diffusion coefficient *D* = 10^−11^ m^2^/s along the height of the channel *h* = 100 μm to be τ_D_ = *h*^2^/(2*D*) = 500 s. These two timescales are much shorter than the timescale of gel contraction (~3 h is Figure 2B) and thus the DNA reaction and kinesin diffusion along the height of the channel do not play a role in the contraction rate of the gel.

### 8. Experiments showing the influence of the invader DNA strand concentration

**Figure S7:**
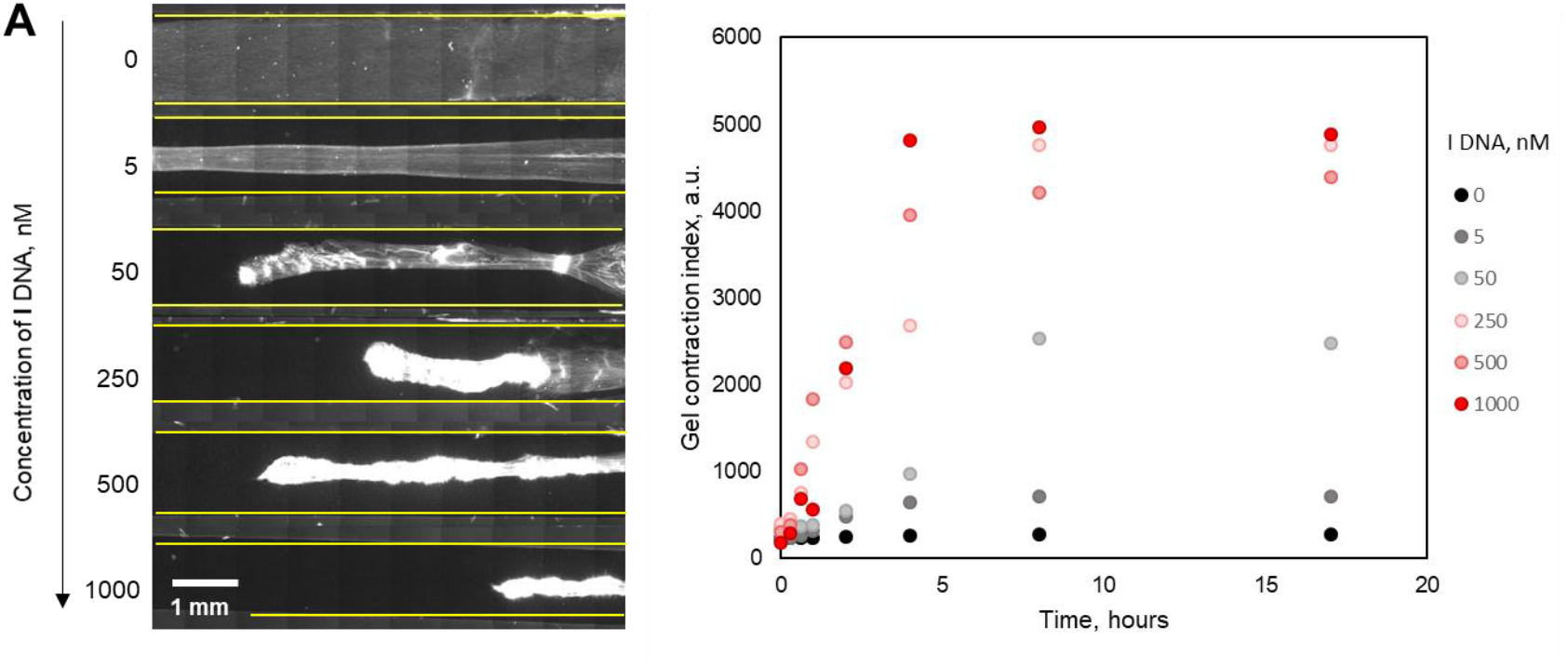
A: Fluorescence microscopy images of the microtubules-containing solution confined in glass channels 10 hours after the addition of the different concentrations of **I** strand. The borders of the channels are marked yellow. B: Corresponding gel contraction index over time. For detailed conditions see Table S2.

### 9. Experiments showing the influence of the initial streptavidin surface density

**Figure S8:**
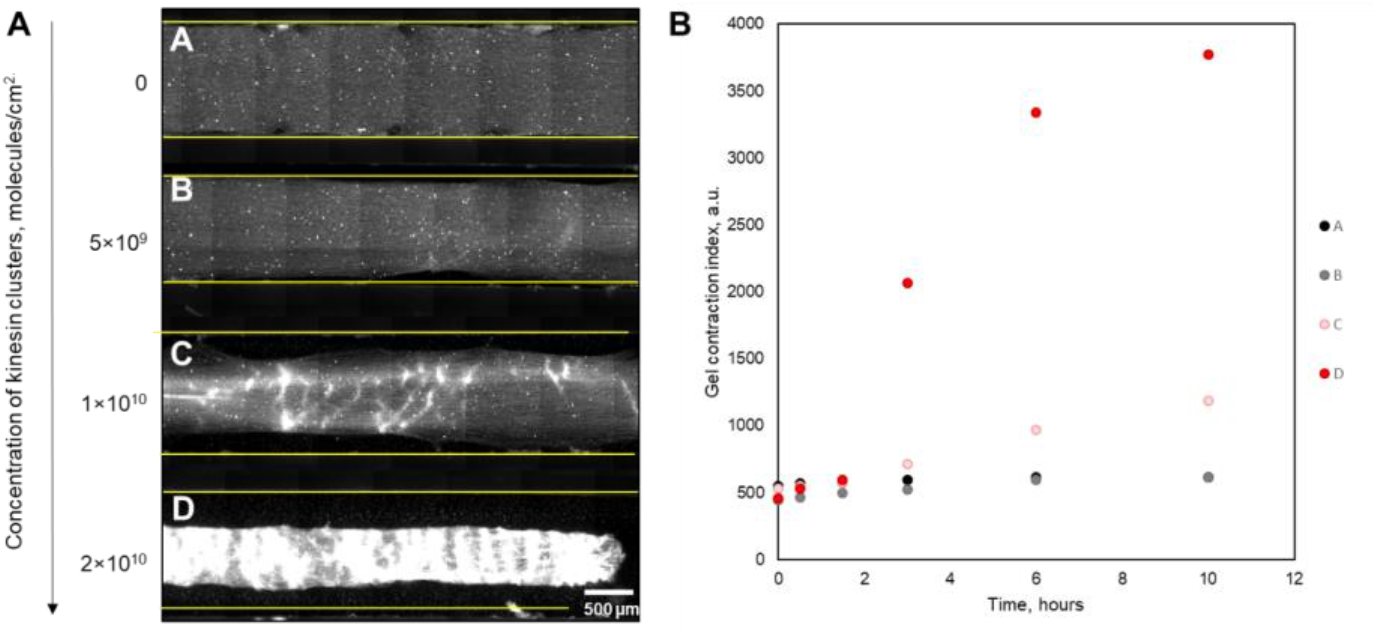
A: Fluorescence microscope images of the microtubule-containing solution confined in glass channels in the absence (A) and in the presence of 1 μM **I** DNA (B, C, D) at different streptavidin surface densities: 5×10^9^ (B), 1×10^10^ (A and C), and 2×10^10^ molecules/cm^2^ (D). The borders of the channels are marked yellow. The images were recorded 17 hours after the addition of **I**. B: Corresponding gel contraction index over time. For detailed conditions see Table S2.

### 10. Experiments showing the diffusion of the invader I DNA through the channel

To illustrate the correlation between diffusing invader DNA and microtubules’ activity, **I** DNA was mixed in 1:1 ratio with **N** DNA that bears fluorescent label to enable diffusion tracking.

**Figure S9:**
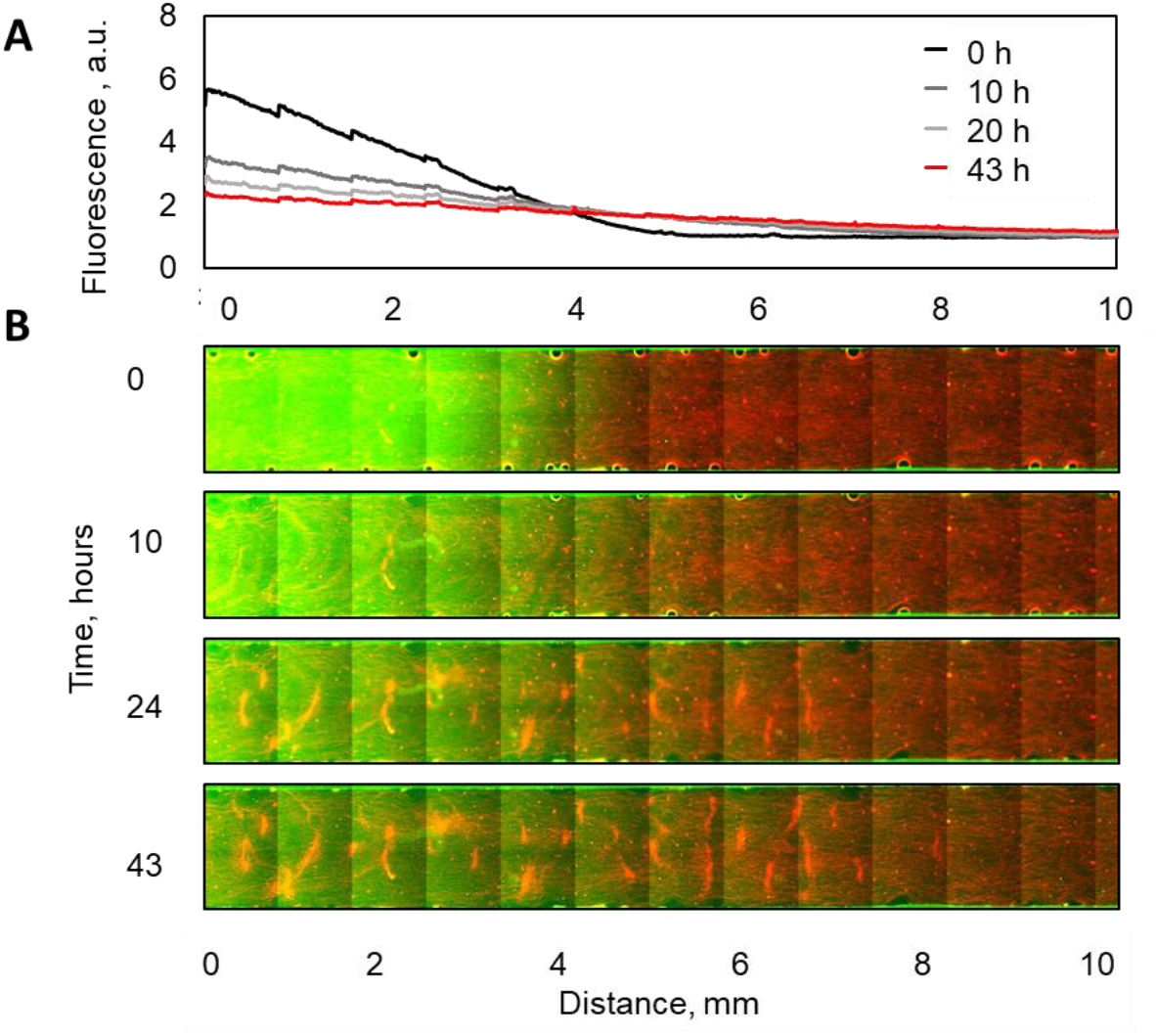
A: The intensity of the fluorescence of the DNA strand **N** used to track the diffusion of **I** DNA strand throughout the channel. B: Time-lapse fluorescent microscopy images of the microtubule containing solution confined in glass channels after the addition of 1 μM of DNAs **I** and **N** on the left side of the channel. Strand **I** activates the gel while inert fluorescent strand **N** helps tracking the diffusion of DNA along the channel. The duration of the experiment is 43 hours. An estimated surface density of the immobilized streptavidin is 1×10^10^ molecules/cm^2^.

### 11. Control experiment showing the importance of the kinesin presence

**Figure S10:**
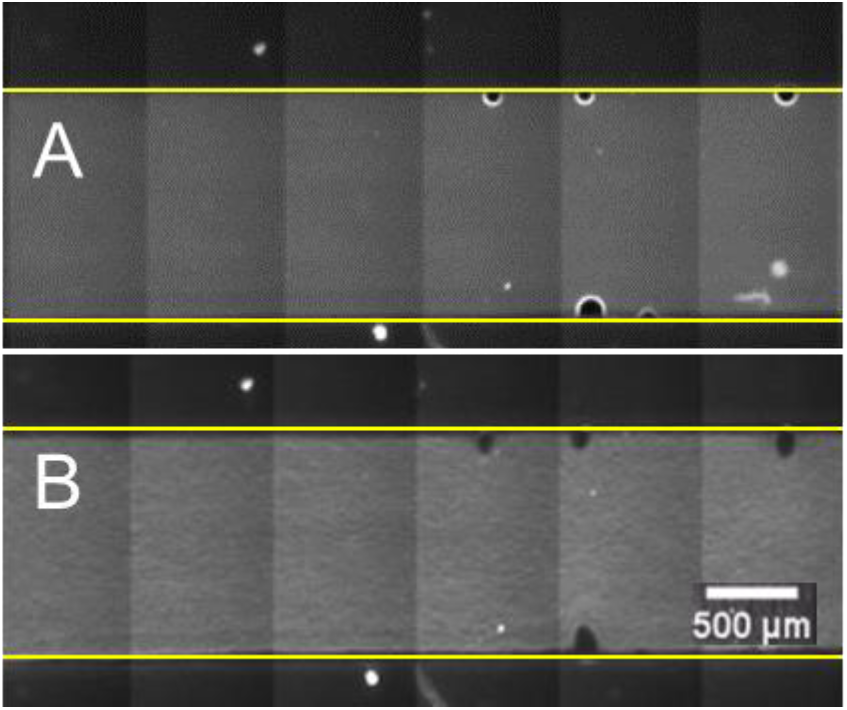
Fluorescence microscopy images of the microtubules-containing solution confined in glass channels before (A) and 10 hours after the addition of 1 μM of **I** strand (B). The borders of the channels are marked yellow. For detailed conditions see Table S2.

### 12. Control experiment showing direct interactions of microtubules with kinesin clusters

In this experiment, we tested the structural changes of the active gel prepared by direct mixing of microtubules with kinesin in solution. Fluorescence microscopy measurements show that the transition from nematic state to corrugated structure appeared earlier compared to system in Figure 1C. These results demonstrate that the interaction between motor protein and microtubule filaments was delayed when DNA strand displacement was involved to trigger the release of the kinesin from the surface. After 90 minutes, the gel started to contract along the x and y axes. The resulting structure exhibits some faint bands but it lacks the periodic bright bands that where clearly visible in Figure 1C. We speculate that slow diffusion of motors from the surface and delayed interaction between microtubules and kinesin might be the reason for bands appearance.

**Figure S11.**
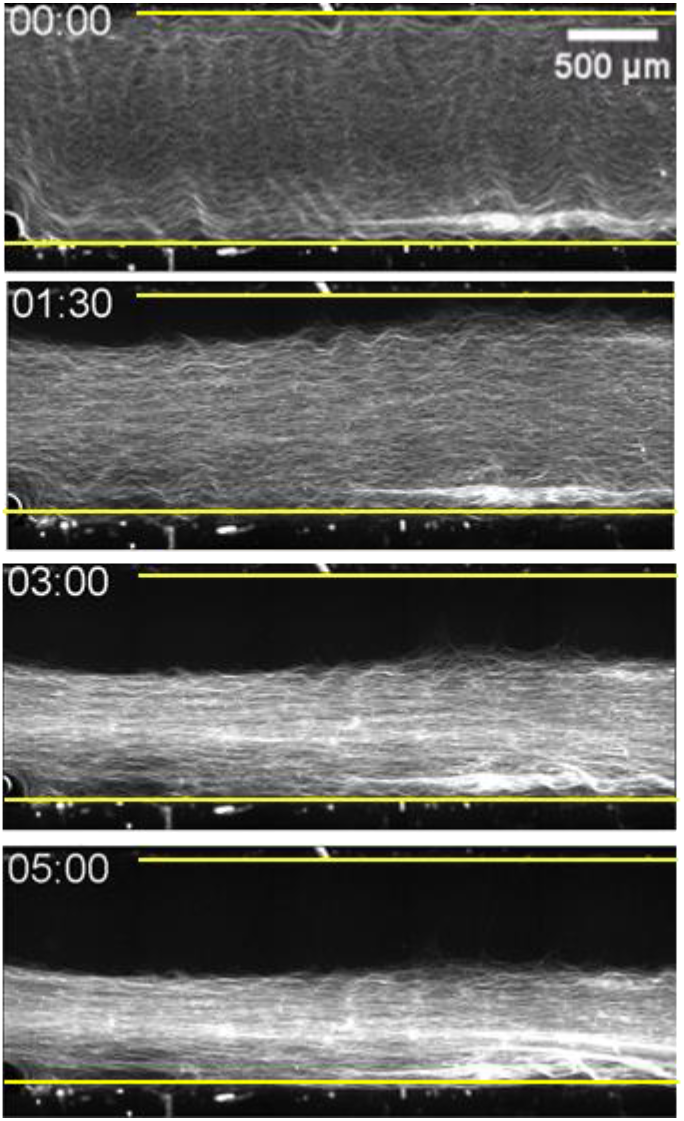
Time-lapse fluorescent microscopy images of the microtubule containing solution confined in glass channels after the addition of kinesin/streptavidin clusters (5 nM/ 5 nM). The duration of the experiment is 5 hours.

### 13. Supplementary Movies

**Movie S1.**
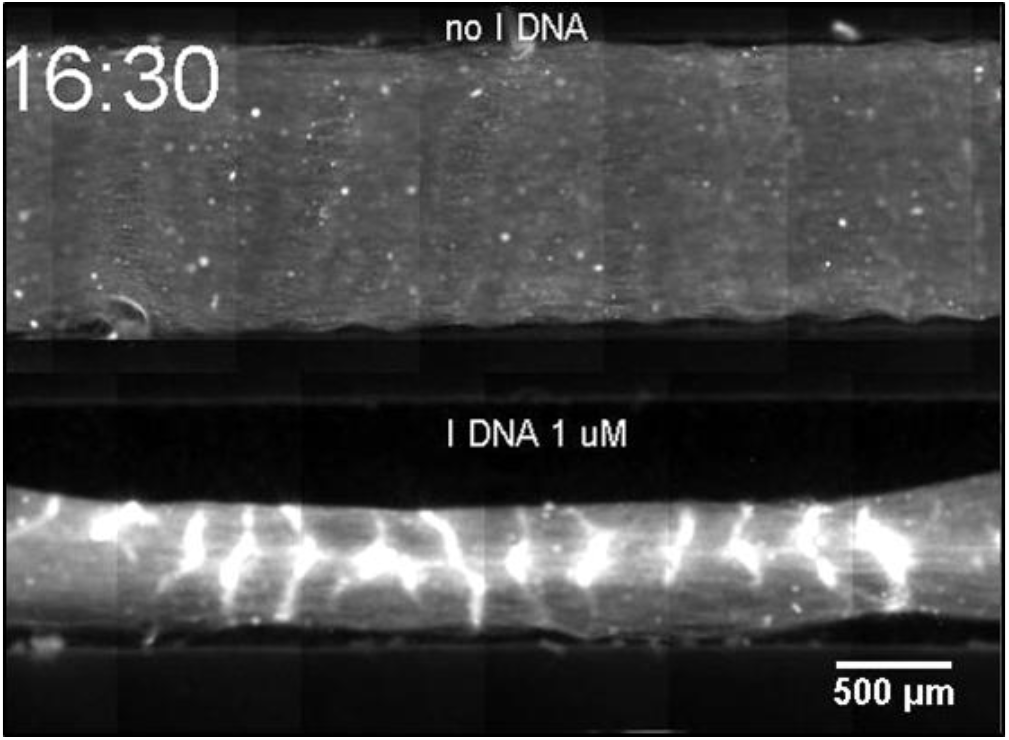
Fluorescent microscopy images of the microtubule solution confined in glass channels in the absence (top) and in the presence of 1 μM of **I** DNA (bottom).

**Movie S2.**
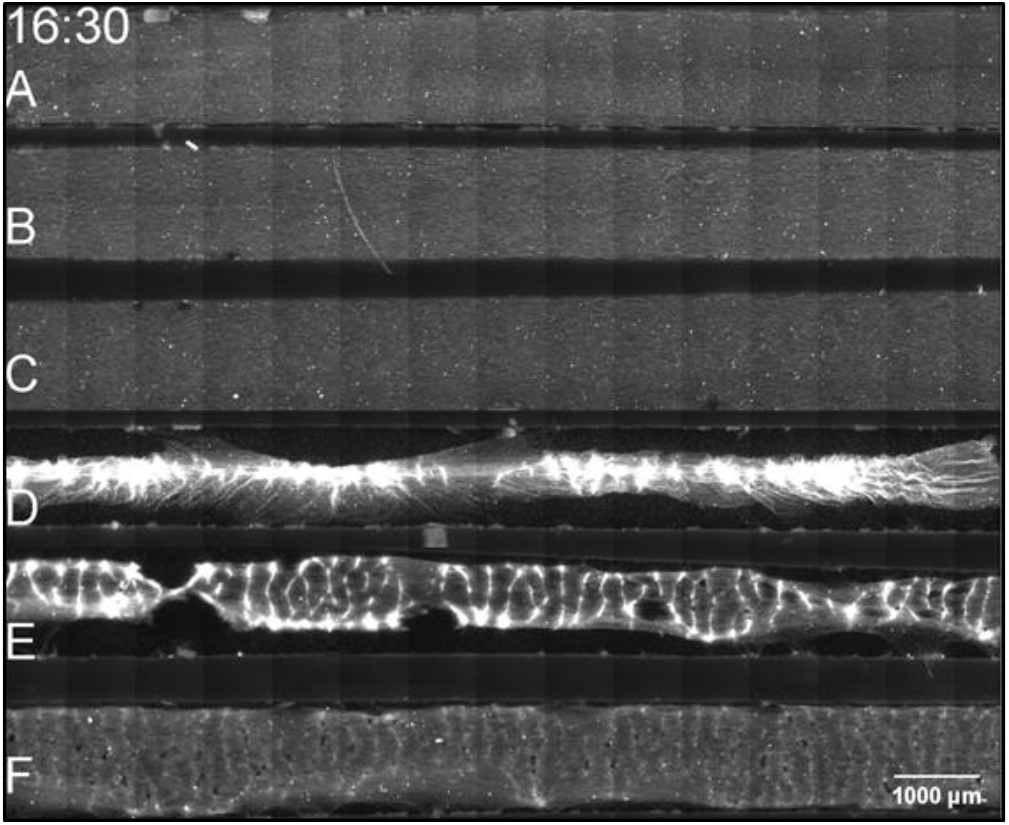
Fluorescence microscopy images of the microtubule solution confined in glass channels modified with biotinylated DNA strands of different complementarity length (15 nt (A and D); 22 nt (B and E); 30 nt (C and F)) in the absence (A-C) and in the presence (D-F) of 1 μM **I** strand. The shadows moving through the images are artifacts due to air bubbles in the oil separating the glass slide and the temperature controlled stage.

**Movie S3.**
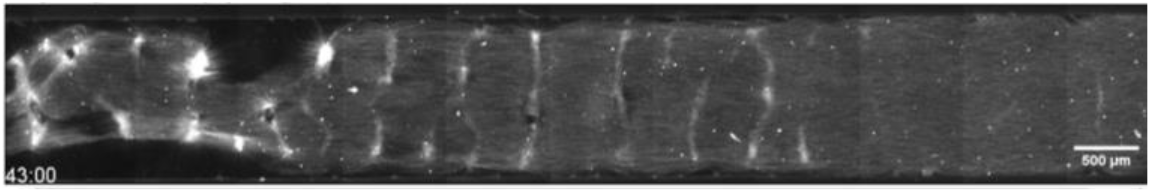
Fluorescence microscopy images of the microtubule solution confined in glass channels after the addition of 0.75 uL of 1 μM **I** DNA on the left-hand side.

**Movie S4.**
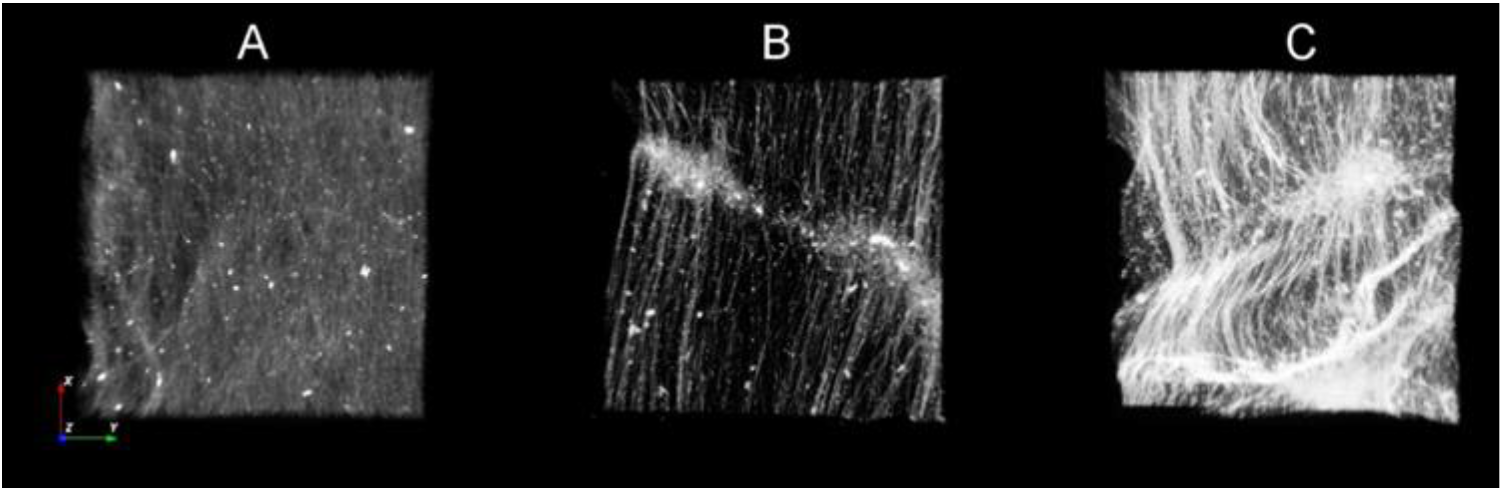
Confocal fluorescence microscopy images of the microtubule solution confined in glass channels modified with biotinylated DNA strands of different complementarity length (30 nt (A and B) and 15 nt C) in the absence of **I** DNA (A), and in the presence of 1 μM **I** strand (B and C).

